# ACPA-IgG variable domain glycosylation increases before the onset of rheumatoid arthritis and stabilizes thereafter; a cross-sectional study encompassing over 1500 samples

**DOI:** 10.1101/2021.11.05.467407

**Authors:** T. Kissel, L. Hafkenscheid, T.J. Wesemael, M. Tamai, S.Y. Kawashiri, A. Kawakami, H.S. El-Gabalawy, D. van Schaardenburg, S. Rantapää-Dahlqvist, M. Wuhrer, A.H.M. van der Helm-van Mil, C.F. Allaart, D. van der Woude, H.U. Scherer, R.E.M. Toes, T.W.J. Huizinga

## Abstract

**Objective:** The autoimmune response in rheumatoid arthritis (RA) is marked by anti-citrullinated protein antibodies (ACPA). A remarkable feature of ACPA-IgG is the abundant expression of *N*-linked glycans in the variable domain. Nonetheless, the presence of ACPA variable domain glycans (VDG) across disease stages and its’ response to therapy is poorly described. To understand its dynamics, we investigated the abundance of ACPA-IgG VDG in 1574 samples from individuals in different clinical disease stages.

**Methods:** Using liquid chromatography, we analyzed ACPA-IgG VDG profiles of 7 different cohorts from Japan, Canada, the Netherlands and Sweden. We assessed 184 healthy, 228 pre-symptomatic, 277 arthralgia, 305 patients at RA-onset and 117 RA-patients 4, 8 and 12 months after disease onset. Additionally, we measured VDG of 234 samples from RA-patients that did or did not achieve long-term drug-free remission (DFR) during up to 16 years follow-up.

**Results:** Our data show that ACPA-IgG VDG significantly increases (p<0.0001) towards disease-onset and associates with ACPA-levels and epitope spreading pre-diagnosis. A slight increase in VDG was observed in established RA and a moderate influence of treatment. Individuals who later achieved DFR displayed reduced ACPA-IgG VDG already at RA-onset.

**Conclusion:** The abundance of ACPA-IgG VDG rises towards RA-onset and correlates with maturation of the ACPA-response. Although, ACPA-IgG VDG levels are rather stable in established disease, a lower degree at RA-onset correlates with DFR. Even though the underlying biological mechanisms are still elusive, our data support the concept that VDG relates to an expansion of the ACPA-response pre-disease and contributes to disease-development.

## Introduction

Rheumatoid arthritis (RA) is a prevalent, slowly evolving autoimmune disease with arthralgia as an important pre-disease manifestation. The most specific autoimmune response for RA is characterized by the presence of anti-citrullinated protein antibodies (ACPA), which can be found several years before the onset of clinical symptoms. ACPA-positive patients have a more severe disease course and a lower chance to achieve drug-free remission (DFR) as compared to seronegative patients [1]. ACPA responses are known to be dynamic during the transition towards RA, as an increase in ACPA levels combined with a broader epitope recognition-profile is associated with the development of clinical symptoms [2]. Autoantibody levels are, however, not associated with long-term treatment response and do not predict DFR [3]. Glycomic analysis revealed that ACPA-IgG are abundantly glycosylated in their antigen-binding fragments expressing complex-type variable domain glycans that are mainly disialylated and bisected [4]. A variable domain glycosylation (VDG) on more than 90% of the autoantibodies, is an outstanding characteristic of ACPA-IgG and distinguishes the molecules from conventional IgG which display a considerably lower VDG of approximately 12% [4, 5]. Although the role and dynamics of ACPA-IgG Fc-glycans has been studied extensively [6-8], little is known about the expression levels or potential biological implications of the ACPA VDG. As carbohydrates might encode important biological information and possibly affect cellular functions, it is important to understand VDG dynamics over time in relation to the disease course. Previously, we have shown that ACPA-IgG VDG can already be present several years before RA-onset. In a Canadian population, ACPA-IgG VDG were predictive for disease development [9, 10]. However, it is still elusive how ACPA-IgG VDG changes between different clinical disease stages from healthy, symptom-free individuals to individuals with arthralgia to patients at RA-onset and with established RA. Additionally, it is unclear whether VDG levels are associated with treatment outcomes, predict DFR and disease flares, or can be modified by treatment. To understand the momentum of VDG and thereby their possible contribution to the autoreactive B-cell responses in RA, we cross-sectionally investigated the presence and abundance of ACPA-IgG VDG in 1574 samples from ethnically diverse individuals in various disease stages. Furthermore, by analyzing samples from a well-controlled treatment strategy trial, the Improved study, aiming to assess the most effective strategy in inducing remission in early RA, we investigated longitudinal VDG changes in established RA after treatment escalation or treatment tapering [11]. Lastly, we longitudinally analyzed ACPA-IgG VDG glycan changes of individuals visiting the Leiden Early Arthritis Clinic (EAC) and achieving drug-free sustained (>1 year) remission (DFSR) or late disease flares [12].

## Materials and Methods

### Study Cohorts

ACPA-IgG VDG were analyzed in 1574 serum samples from individuals in different clinical disease stages. The descriptive cohort data are presented in tables S1 and S2. Additionally, 247 healthy donor control samples, 150 ACPA-positive RA and 101 ACPA-negative RA control samples were assessed.

#### Cohort 1, healthy symptom-free (Japan, Nagasaki)

Healthy symptom-free individuals (n=58) were included that were tested positive for the presence of ACPA and are part of the Nagasaki Island Study performed in Japan (a prospective cohort study based on resident health check-ups with a follow-up of three years) [13]. 9 individuals (15.5%) developed RA during follow-up.

#### Cohort 2, healthy and RA- onset (Canada, Manitoba)

Samples of ACPA-positive healthy individuals (n=126) (first degree relatives of RA-patients) and 23 paired samples at RA- onset (n=23) were included. Individuals were part of the longitudinal research project ‘Early Identification of Rheumatoid Arthritis in First Nations’ based at the Arthritis Centre at the University of Manitoba [14].

#### Cohort 3, pre-symptomatic and RA- onset (Sweden, Umea)

ndividuals, diagnosed with RA later in life, were retrospectively selected prior to symptom-onset (n=228, median (IQR) pre-dating time: 4.7 (5.9) years) and after diagnosis of RA (n=126). The pre-disease samples were derived from the Medical Biobank of Northern Sweden [15].

#### Cohort 4, arthralgia (the Netherlands, Amsterdam, Reade)

ACPA-positive individuals with arthralgia (n=239) prospectively sampled at rheumatology outpatient clinics in the Amsterdam area of the Netherlands [16] were selected. Individuals were followed up to 10 years and 137 (57.3%) developed arthritis during follow-up.

#### Cohort 5, arthralgia (the Netherlands, Leiden, CSA)

Individuals with arthralgia (n=38) and paired RA- onset samples (n=26) were included. Individuals, at risk of RA development, were recruited for the prospective Clinically Suspect Arthralgia (CSA) cohort in the Leiden Rheumatology outpatient clinic and followed longitudinally [17].

#### Cohort 6, RA- onset and established RA (the Netherlands, Leiden, Improved)

Longitudinal samples of 130 RA patients at disease-onset, 4 months (n=117), 8 months (n-112) and 12 months (n=117) were included. Individuals, recruited in 12 hospitals in the western area of the Netherlands, were included in the Induction therapy with Methotrexate and Prednisone in Rheumatoid Or Very Early arthritic Disease (Improved) study. This multicenter, randomized control trial was aimed to achieve DFR including treatment alteration every 4 months. Initial treatment was methotrexate (MTX) and prednisone, followed by either tapering of medication (early remission) or randomization to one of two treatment arms: MTX, prednisone, hydroxychloroquine, and sulfasalazine combination (arm 1) or MTX and adalimumab combination (arm 2) [11]. At 8 months patients tapered and discontinued methotrexate, if they achieved early remission.

#### Cohort 7, RA- onset, DFR, DFSR and late disease flares (the Netherlands, Leiden, EAC)

Individuals at RA- onset that did not achieve DFR at later time-points (n=59) and longitudinal samples (n=175) of individuals that achieved DFR (n=41) or sustained DFR (DFSR) (n=35) were selected and recruited from the Leiden Early Arthritis Clinic (EAC) in the Leiden University Medical Center (the Netherlands) [12]. Samples at RA- onset, the pre-remission phase, DFR, DFSR and if present at the time point of late disease flares were included with a follow-up of up to 16 years.

### ACPA-IgG capturing and VDG analysis using liquid chromatography

Capturing of ACPA-IgG, total glycan release, glycan labeling and purification was performed as previously described [9]. In brief, ACPA were affinity isolated from 25µl serum samples using NeutrAvidin Plus resin (Thermo Scientific) coupled with 0.1µg/µl CCP2-biotin followed by IgG capturing using FcXL affinity beads (Thermo Scientific). *N*-linked glycans were released using PNGaseF, subsequently labelled with 2-AA and 2-PB and HILIC SPE purified using GHP membrane filter plates (Pall Life Science). Ultra-high performance liquid chromatography (UHPLC) was performed on a Dionex Ultimate 3000 (Thermo Fisher Scientific) instrument, a FLR fluorescence detector set and an Acquity BEH Glycan column (Waters, Milford, MA). Separation and glycan peak alignment were performed as previously published [9]. HappyTools version 0.0.2 was used for calibration and peak integration [18]. The *N*-linked glycan abundance in each peak was expressed as the total integrated area under the curve (AUC). The cut- off was based on blank and ACPA-negative healthy donor samples, excluding outliers (below or above Q_1_−1.5 x IQR). The percentage of ACPA-IgG VDG was calculated based on the following formula: [(G2FBS1+G2FS2+G2FBS2)/(G0F+G1F+G2F) x100] (figure S1) [19].

### Statistical analyses

Continuous data were analyzed using non-parametric methods (Kruskal-Wallis test for non-paired samples including multiple comparisons and Mann-Whitney’s U-test for non-paired samples) and parametric tests (Mixed-effect analysis for matched-paired samples including missing values) when appropriate. Correlations between ACPA levels (log transformed) and percentages of VDG were assessed with Pearson correlation. All p-values are two-sided and p<0.05 was considered as statistically significant. Logistic and ordinal regression analyses were performed for cohort 4 (arthralgia, Amsterdam) and cohort 6 (RA-onset, Leiden) to investigate the association of ACPA-IgG VDG/ACPA levels with epitope spreading, remission and early DFR. The unstandardized coefficient (B) represents the mean change in the response given a one unit change in the predictor. The longitudinal and repeated measures data from cohort 6 (RA-onset and established RA, Leiden) were analyzed using generalized estimating equations (GEE), as specified before [3].GEE was used to assess VDG changes over time and associations with treatment/treatment decisions. The specific covariates and dependent variables are listed in the table legends, respectively. Statistical calculations were performed using STATA (V.16.1; STATA Corp, College Station, Texas USA).

## Results

### ACPA-IgG variable domain glycosylation increases towards disease onset and remains stable in established RA

To provide a comprehensive overview of the presence and abundance of ACPA-IgG VDG (figure S1a and b) we analyzed 1450 ACPA-positive and 124 ACPA-negative samples from individuals in different clinical disease stages (figure 1a and table S1). Comparable with previous studies [9, 10], we identified high percentages of VDG (median of 58.1%) on ACPA-IgG already in healthy individuals (n=184) without symptoms (figure 1b and c). Our cross-sectional analysis revealed a significant increase in VDG (median of 74.7%) in clinically identified individuals with imminent RA (arthralgia) (n=277) compared to healthy individuals (figure 1b, c and S3). An additional significant rise in VDG of 18% was observed when individuals were sampled at RA- onset (n=305, VDG median of 92.6%) (figure 1b, c and tableS1). In established RA (n=346), ACPA-IgG VDG remained stable, with only a moderate increase after 12 months to a median of 105.2% (figure 1b). As previously shown [9], an increase in ACPA-IgG VDG towards RA-onset was also observed in a Swedish population of ACPA- positive individuals that later developed RA. The extended dataset used here also depicts a rise in VDG when analyzed per individual in a longitudinal manner [20], however no significance could be detected cross-sectionally (figure S1d).

**Figure 1:**
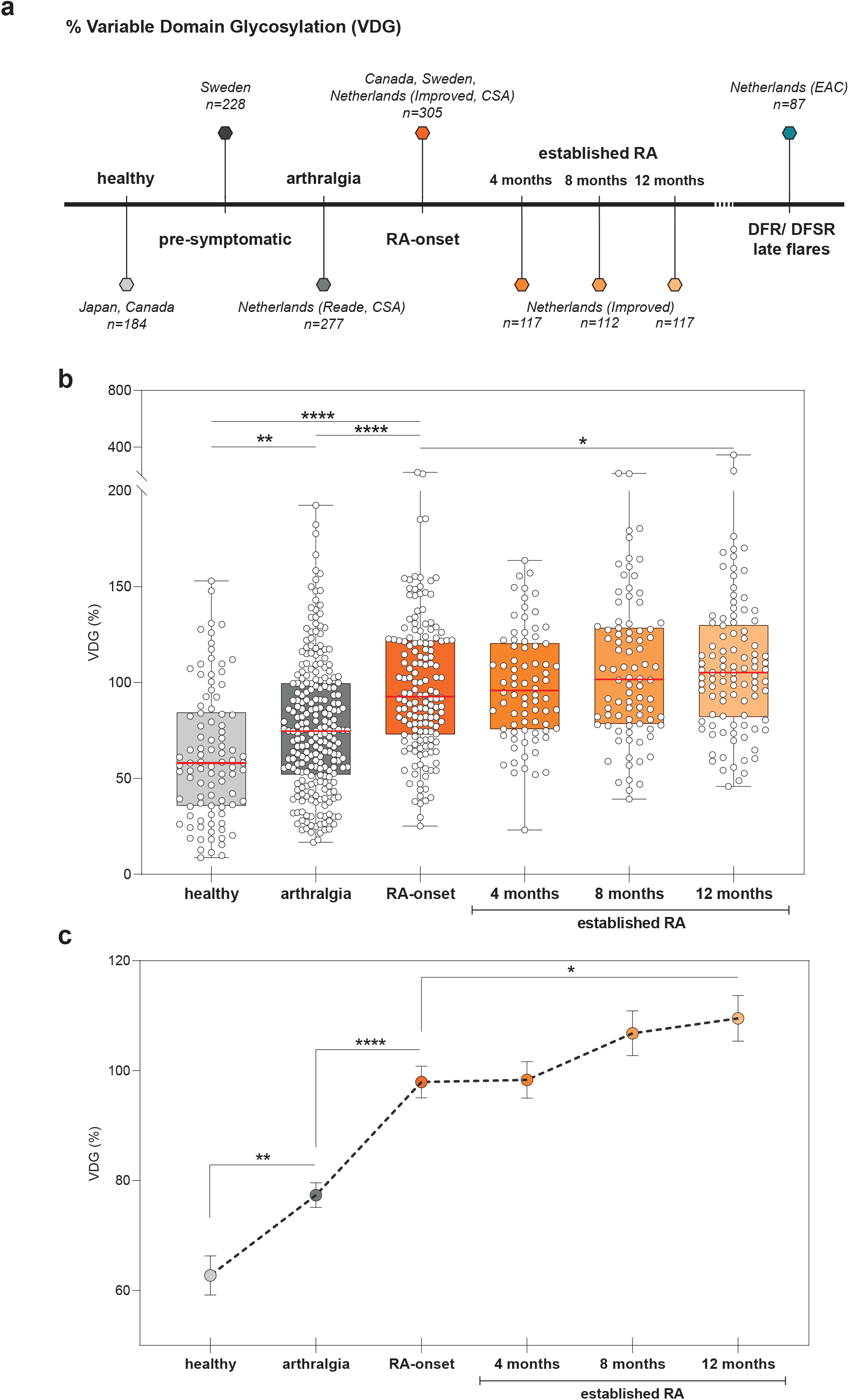
Percentage ACPA-IgG variable domain glycosylation (VDG) of individuals in different clinical disease stages of RA. (a) Schematic ‘time-line’ of different clinical disease stages, the corresponding analyzed cohorts and absolute numbers of analyzed samples. (b) Percentage ACPA-IgG VDG was measured by liquid chromatography and cross-sectional and longitudinal data of healthy, arthralgia, RA-onset and established RA (4, 8 and 12 months) individuals specified in a are shown. Data are presented as box and whiskers including all data points. VDG increased significantly towards disease onset and an additional slight increase 12 months after RA- onset was observed. (c) Percentage ACPA-IgG VDG mean including standard error of the mean (SEM) of healthy, arthralgia, RA-onset and established RA (4, 8 and 12 months) individuals specified in a. Increase in VDG towards disease onset is depicted by the dotted line. Kruskal-Wallis tests were performed for cross-sectional, non-parametric non-paired samples. Mixed-effects analysis was performed for longitudinal, parametric matched- paired samples including missing values. Significant differences are denoted by *(p=0.0387), **(p=0.0026), or ****(p<0.0001).

Overall, the results obtained indicate that the presence of VDG on ACPA-IgG is low in healthy individuals, but increases in individuals that later develop RA. However, in established disease, no further progression of ACPA-VDG is observed in this cross-sectional setting.

### The increase in variable domain glycosylation and maturation of the ACPA immune response are interconnected

To obtain more insights into ACPA-IgG VDG, we investigated the possible association between VDG percentages and the “ maturation” of the ACPA- response by analyzing ACPA levels and the broadness of the citrullinated epitope-recognition profile. Pearson correlation analyses depicted a strong, highly significant correlation between VDG percentages and ACPA levels in healthy individuals (r=0.728 and r=0.672) and for subjects with arthralgia (r=0.640) (figure 2a and S1b). At RA-onset (r=0.131 and r=0.214) and in established RA (r=0.341, r=0.362 and r=0.215), however, we observed only moderate correlations as depicted by the correlation coefficient r (figure 2a and S2). Likewise, our data revealed that ACPA-IgG with increased VDG showed a significantly broader recognition profile towards multiple citrullinated epitopes (figure 2b and c). Ordinal regression analyses confirmed these findings for individuals with arthralgia (p<0.001) (table S3) as well as for patients at RA- onset (p=0.004) and over time in established RA (p<0.001) (table S4)

**Figure 2:**
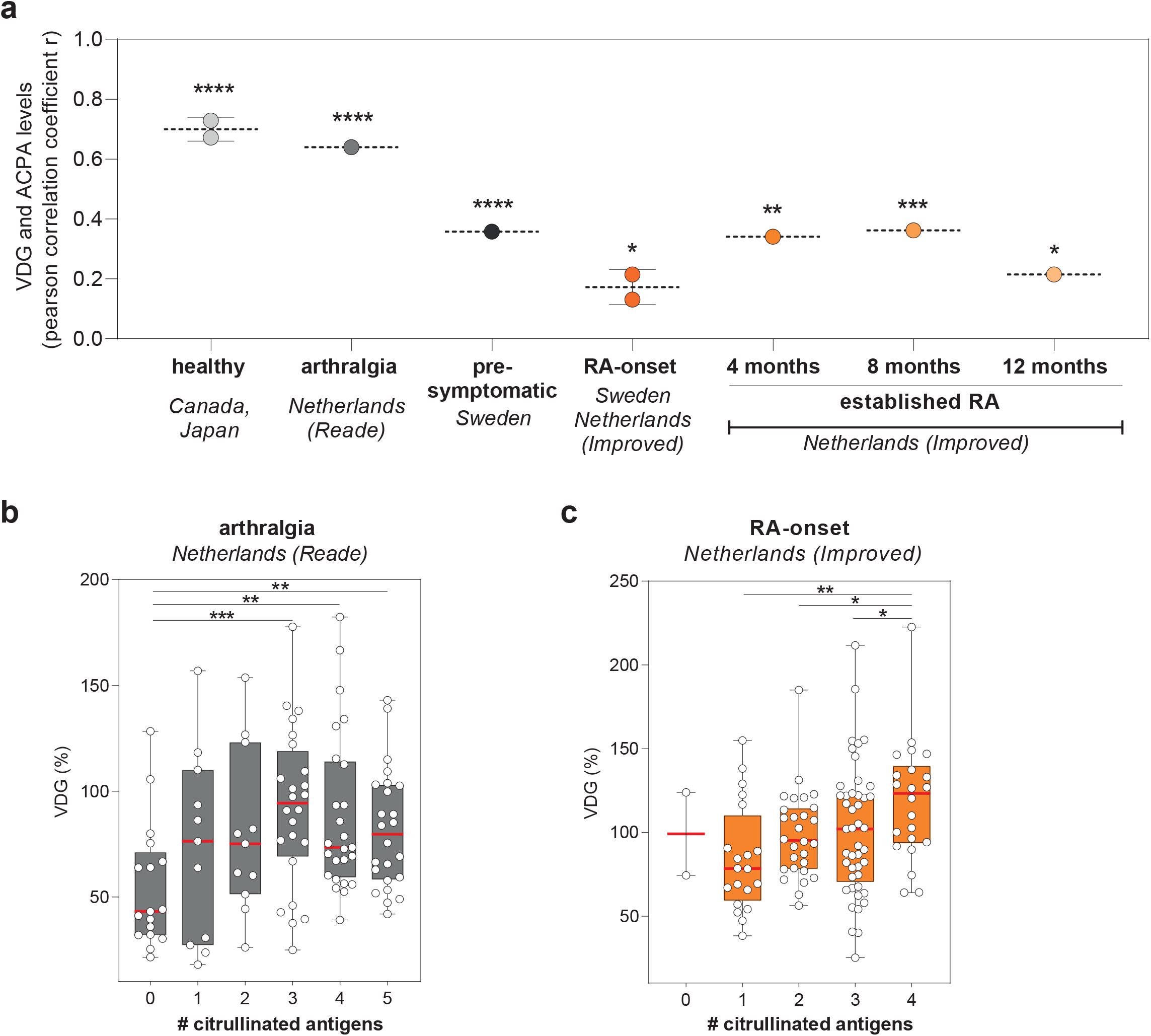
ACPA-IgG VDG correlate with ACPA levels and epitope spreading (maturation of the ACPA response). (a) Pearson correlation coefficients r for correlation between ACPA-IgG VDG and ACPA levels. A strong correlation was observed in the pre-disease phase and a moderate correlation in disease. P-values are two-tailed and significant differences are denoted by *(RA-onset) p=0.0203 (Netherlands), p=0.162 (Sweden), *(12 months) p=0.0338, ** p=0.0023, *** p=0.0006 or **** p<0.0001. (b) VDG percentages shown for ACPA-IgG isolated from arthralgia individuals (*Netherlands, Reade*) and tested for binding to 0-5 different citrullinated antigens. A significant increase in epitope spreading (recognition of cit-antigens) was observed with increasing VDG percentages. Kruskal-Wallis tests were performed for non-parametric non-paired sample: **(0 vs. 4) p=0.0036, **(0 vs. 5) p=0.0096, *** p=0.0006. (c) VDG percentages shown for ACPA-IgG isolated from individuals at RA-onset (*Netherlands, Improved*) and tested for recognition of 0-4 different cit-antigens. Significant increase in recognition of cit-antigens was observed with increasing VDG percentages. Kruskal-Wallis tests were performed for non-parametric non-paired sample: **(2 vs. 4) p=0.0411, **(3 vs. 4) p=0.0417, *** p=0.0011.

Thus ACPA-IgG VDG associates with ACPA levels and the breadth of the epitope-recognition profile, suggesting that these two features of the ACPA-responses are interconnected.

### The impact of immunosuppression on ACPA-IgG variable domain glycosylation

By taking advantage of the design of the Improved study (figure 3c), we investigated whether ACPA-IgG VDG predict early remission or associate with the intensity of immunosuppression. First, we used the longitudinal data set to identify ACPA-IgG VDG changes over time by analyzing matched-paired individuals at RA- onset (n=130) versus 4 (n=117),8 (n=112) and 12 months (n=117) after disease development. VDG appeared to be steadily and abundantly expressed on ACPA-IgG in disease, although minor changes in expression levels were observed in time. A slight, but non-significant dip was observed 4 months after disease-onset and initiation of methotrexate (MTX) and prednisone treatment (figure 3a,b and table S5). Previous studies have shown this drop upon treatment also for ACPA levels pointing again towards the correlation between VDG and ACPA levels [3]. After 4 months, medication was tapered when patients achieved early remission (MTX only) or individuals were randomized to one of two treatment escalation arms (arm 1: MTX, prednisone, hydroxychloroquine, and sulfasalazine combination, arm 2: MTX and adalimumab combination) (figure 3c) [11]. At 8 months individuals in the early remission group either continued MTX treatment combined with prednisone (no drug-free) or medication was tapered (drug-free), if they were still in remission. Individuals from the treatment escalation group (arm 1 and arm 2) continued MTX in combination with an adalimumab treatment. Overall, irrespective of the treatment arm, VDG increased moderately but significantly 12 months after RA-onset (p=0.037) (figure 3a,b, S4b and table S5). When comparing the different treatment groups, small, but statistically significant impacts of immunosuppression on ACPA-IgG VDG can be observed 12 months after RA-onset (figure 3d and S4a), although absent at 4 and 8 months. This moderate, but significant negative impact of immunosuppression on VDG was confirmed by a generalized estimating equation (GEE) analysis over time (8 vs. 12 months) (DFR: 12.27 (−7.32-31.87) vs. treatment escalation: 6.42 (−0.35-13.10); p=0.007) (table S6) and also observed for ACPA levels in a previous study [3]. Lastly, we investigated, if VDG percentages at RA-onset predict remission after 4 months and early drug-free remission within the first year. Similar to ACPA levels [3], VDG percentages did not predict early (drug-free) remission (table S7).

**Figure 3:**
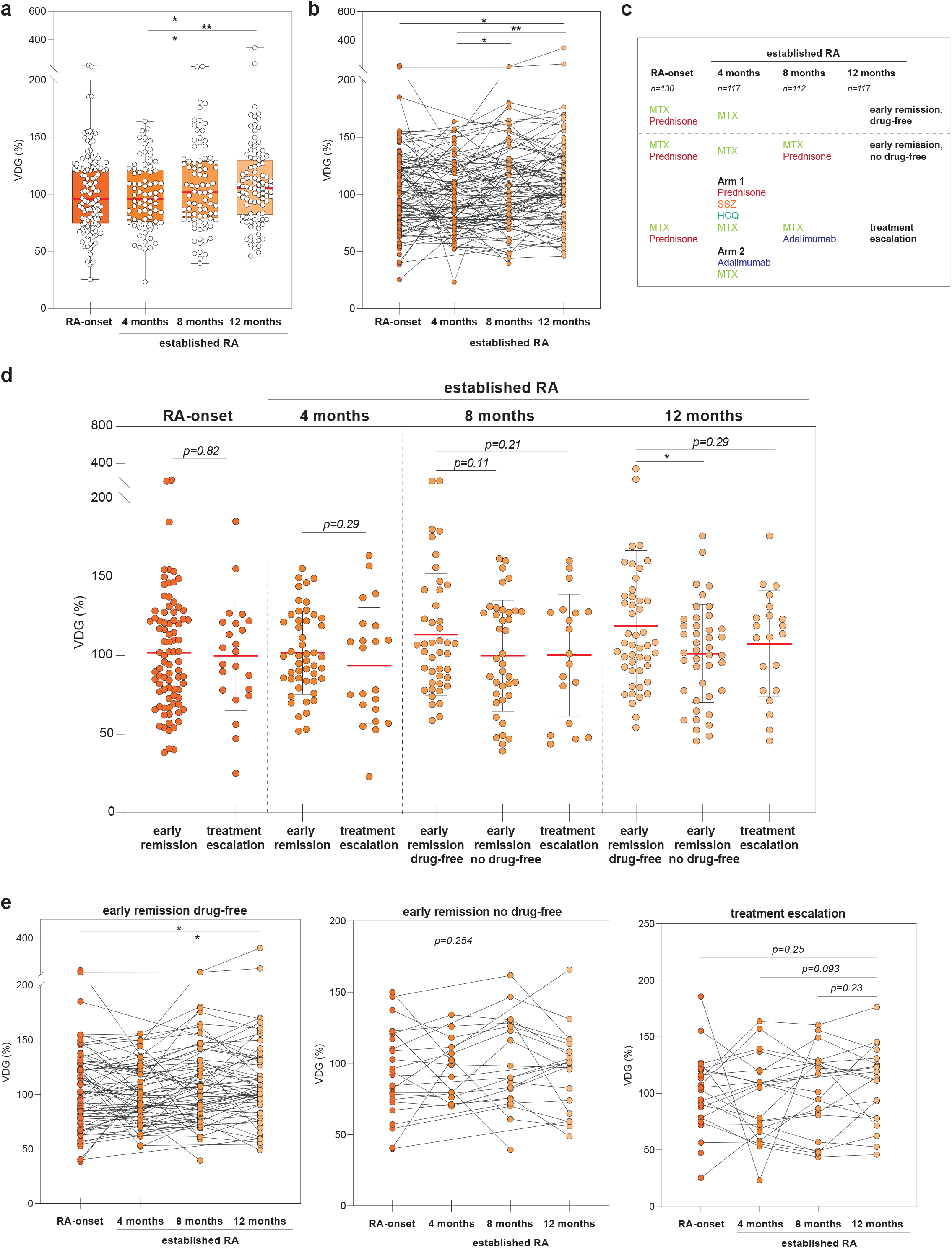
Longitudinal analysis of ACPA-IgG VDG at RA- onset and in established RA (*Netherlands, Improved*). ACPA-IgG VDG showed a non-significant dip 4 months and a slight, but significant increase 12 months after RA-onset. (a) Data are presented as box and whiskers including all data points. *(RA-onset vs. 12 months) p=0.0242, *(4 vs. 8 months) p=0.0373, **p=0.0024. (b) Matched paired longitudinal samples are presented. *(RA-onset vs. 12 months) p=0.0242, *(4 vs. 8 months) p=0.0373, **p=0.0024. (c) Schematic illustration of the treatment groups. HCQ=hydroxychloroquine, MTX=methotrexate, SSZ=sulfasalazine (d) ACPA-IgG VDG percentages shown for treatment groups: early remission (drug-free), early remission (no drug-free) and treatment escalation. The data showed a drop in the no drug-free early remission and treatment escalation group compared to the drug-free early remission group after 12 months. Ordinary one-way ANOVA for parametric non-matched samples: *p=0.026. (d) Longitudinal matched paired samples are shown for the early remission (drug-free and no drug-free) and treatment escalation groups. The data showed a slight increase in VDG 12 months after RA-onset and a marginal dip after 4 months. *(RA-onset vs. 12 months p=0.0352, *(4 vs. 12 months) p=0.0088. Mixed-effects analysis was performed for parametric matched-paired samples including missing values.

### Individuals achieving sustained DFR show decreased variable domain glycosylation in active disease

As a next step, we performed cross-sectional and longitudinal ACPA-IgG VDG analyses of individuals that achieved long-term DFSR or experienced DFR with late flares. We therefore made use of the unique EAC database including patients that were followed up to 18 years after disease-onset. Using this database, we were able to identify and approach 41 individuals that had achieved DFR and 35 patients achieving long-term (>1 year) DFR. The longitudinal analysis was performed for matched-paired patients at RA- onset (n=36), in active disease (pre-remission) (n=52), at the time point of DFR (n=41), DFSR (>1 year) (n=35) and when experiencing late disease flares (n=11). Again, the data show that VDG are stably expressed in disease. Intriguingly, however, patients that achieved DFR during follow-up (n=36, VDG=57.9%) showed significantly reduced ACPA-IgG VDG at the onset of disease compared to age and gender matched patients that did not achieve remission and presented a persistently high disease activity score (DAS>3) (n=59, VDG=83.8%) (figure 4a and table S2). In contrast, no statistically significant changes were observed when the ACPA-IgG VDG percentages were determined over time in the DFR- or any of the other groups analyzed (figure 4b-d).

**Figure 4:**
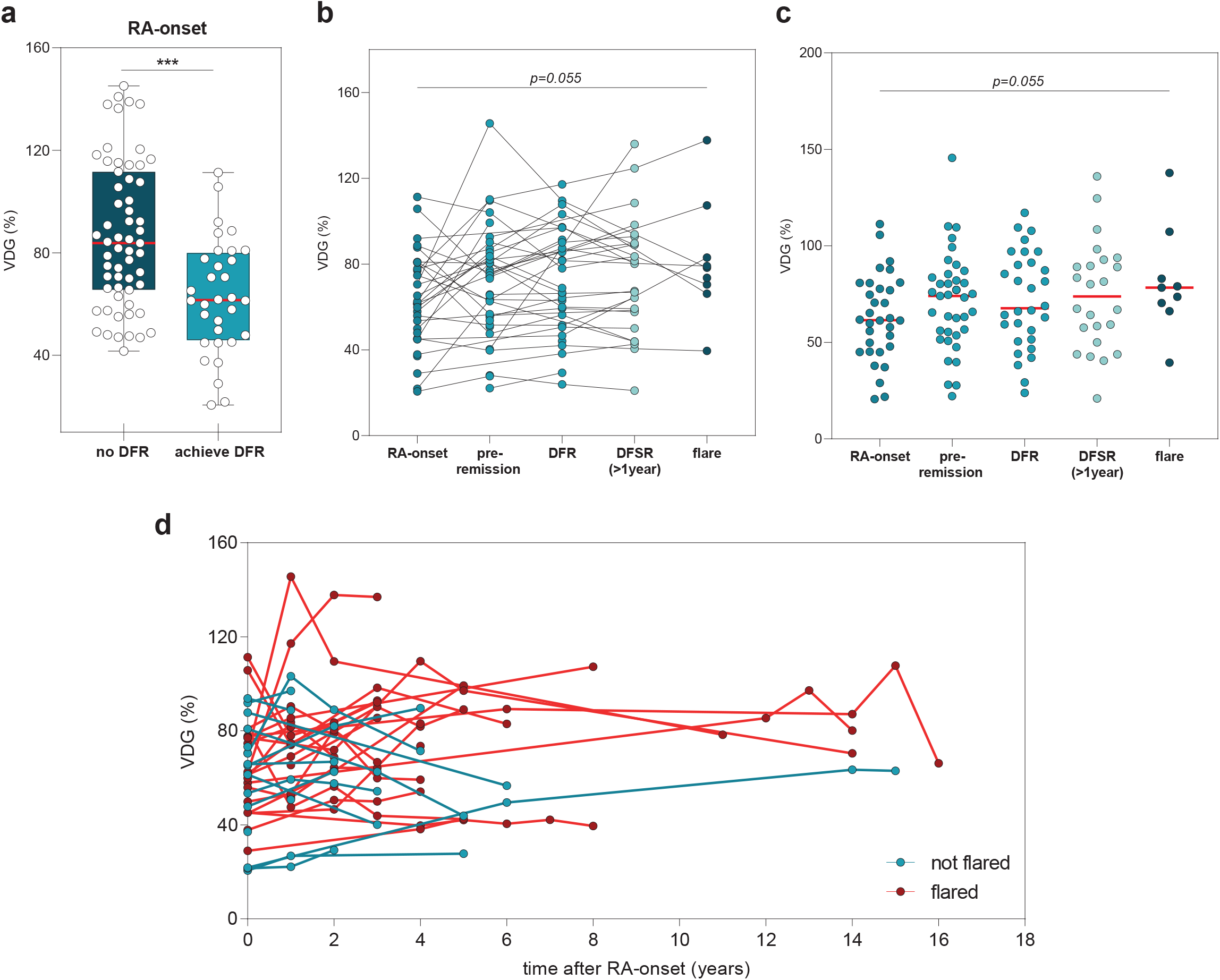
Cross-sectional and longitudinal analysis of ACPA-IgG VDG at RA- onset and in drug-free remission (DFR) (*Netherlands, EAC*). (a) ACPA-IgG VDG percentages at RA- onset for individuals that don’t achieve remission compared to individuals that achieve DFR. Individuals that develop DFR later in time showed significantly reduced VDG percentages at disease onset. (b) ACPA-IgG VDG of matched-paired samples that achieve DFR, sustained DFR or DFR with late flares. VDG percentages increased non-significantly towards DRF, showed a marginal drop in DFR and were slightly higher in individuals that flare. (c) Same individuals as in b are shown as scatter dot plots with the mean of each group indicated as red line. (d) Time-line of longitudinal EAC samples. Individuals that flare later in time are depicted in red and individuals that stay in DFR are depicted in turquoise. Individuals that flare showed a slight increase in VDG over time, while individuals that stayed in DFR marginally decreased in VDG. Mann-Whitney test was performed for non-parametric, non-matched samples: ****p<0.0001. Mixed-effects analysis was performed for parametric matched-paired samples including missing values.

Thus, this longitudinal time-line, confirms that ACPA-IgG express a constant amount of VDG after RA- onset. The cross-sectional dataset also indicates that individuals that achieve long-term DFR have introduced less glycans into the variable domains of their ACPA-IgG at RA- onset.

## Discussion

An important key characteristic of IgG autoantibodies from RA- patients is the abundant presence of bisected and disialylated glycans in the variable domain. To gain insight into the introduction and occurrence of this unusual antibody feature across different disease stages, we have captured ACPA-IgG of more than 1500 samples from 930 individuals in different clinical disease stages. Moreover, we have analyzed the effect of therapy on the degree of variable domain glycosylation on ACPA. The high sample size increases the power of our study and we demonstrate that ACPA-IgG VDG correlates strongly with the maturation of the ACPA immune response pre-disease. We show that the abundance of ACPA-IgG VDG increases significantly from healthy ACPA-positive individuals (58.1%) towards the pre-RA phase (arthralgia) (74.7%) with a further increase towards disease-onset (92.6%). In established RA, we noted a constant high expression of glycans on the variable domain of ACPA-IgG with a slight, but significant, increase after 12 months (105.2%). These latter findings are in agreement with our previous observations, estimating more than 90% VDG on ACPA-IgG in RA [4] as well as the finding that >80% of ACPA B cell receptors in RA express *N*-linked glycosylation- sites in the variable region [21]. Increased VDG levels were mainly observed in individuals who tapered treatment, while patients that experienced more intense treatment show reduced ACPA-IgG VDG profiles in time. This significant impact of immunosuppression was also observed for ACPA levels [22], confirming the correlation between ACPA levels and ACPA VDG, which we most strongly observed in the pre-disease phase. These findings are also in line with the notion that VDG could have a regulatory impact on the ACPA immune response. In this respect, it is intriguing to note that the HLA-Shared Epitope alleles predispose to ACPA harboring VDG [9], also after correction for ACPA levels, and thus linking ACPA VDG with the major genetic risk factor for RA. Of note, we observed that individuals achieving long-term drug-free remission present lower VDG profiles in active disease (58%) compared to patients that did not reach long-term remission (84%). The relevance of these findings are unknown, although it is remarkable that long-term DFR, a relatively rare event in ACPA-positive RA, connects to a lower VDG on ACPA.

Importantly, reduced ACPA levels are not the cause of a lower variable domain glycosylation, which was controlled by titrating ACPA-IgG into healthy serum samples resulting in a maintained high degree of VDG (figure S5b). Thus it is tempting to speculate that VDG serve as an additional ‘hit’ determining the fate of the autoreactive B cell response and thereby impact on ACPA levels.

Together with previous data, showing that *N*-linked glycan sites are selectively introduced into the ACPA B cell receptor sequences upon somatic hypermutation [21] and that VDG are significantly elevated in ACPA-positive individuals that transition to disease [19], our data point towards the hypothesis that a glycan attached to the variable domain is fostering a breach of tolerance of autoreactive B cells. As carbohydrates are known to affect cellular functions, ACPA expressing B cells may gain a selection advantage when abundantly expressing glycans in their variable domains. The disialylated, and thus negatively charged glycans attached to the variable domain, which have also a large steric requirement, might modulate binding to autoantigens or affect B cell receptor signaling of the citrullinated antigen-directed B cells. Further, it cannot be ruled out that VDG impact on effector mechanism and thereby autoantibody-mediated inflammation, similar to Fc-glycans. Next to these areas for further research, it would be interesting to investigate changes in specific VDG traits in more depth as an altered glycan compositions could be associated with defined biological implications as also observed for Fc- glycans. Recent publications show for example that not only Fc-glycans on total IgG, but also ACPA-IgG VDG show a decrease in the bisecting GlcNAc after COVID-19 infection [23, 24].

A limitation of our study is that VDG profiles could only be detected for 70% of the samples analyzed, mainly explained by limited sample amounts or low ACPA levels, as observed in the group of healthy individuals. Especially for ‘rare’ disease stages, such as for the ‘DFR with late flares’ group, only a limited number of samples were available to us. In addition, our conclusions are based on a cross-sectional study as no longitudinal data across all the different clinical disease stages were available. Consequently, the trajectory of individual patients could be different and VDG possibly more stable throughout the disease phases. For example in a cross-sectional setting it cannot be excluded that ACPA-IgG VDG were already increased before disease development in individuals at high risk to develop RA. Importantly however, we did observe an increase in ACPA-IgG VDG towards RA- onset within a previously analyzed longitudinal dataset of pre-symptomatic individuals over a time period of 15 years [20].

In summary, we provide a comprehensive overview of the expression of VDG on ACPA-IgG over various clinical disease stages in RA. Although the biological implications of VDG attached to antibodies in general and ACPA specifically are still largely unexplored, our data show that VDG are a key characteristic of ACPA found in individuals that will develop RA from different ethnicities and across disease stages. Our results show an increase in VDG towards disease progression and suggest, together with previous data indicating a selective introduction of these *N*-linked glycan sites, that VDG may serve as a trigger for the maturation of the ACPA immune response. Therefore it will be relevant to understand the biological impact of VDG on the ACPA immune response and its detailed clinical implications.

## Acknowledgements

The authors would like to thank the Department of Biobank Research at Umeå University, Västerbotten Intervention Programme, the Northern Sweden MONICA study and the County Council of Västerbotten for providing data and samples. We would like to thank Dr. Jan Wouter Drijfhout (LUMC, Leiden) for providing the CCP2 peptide and Carolien Koeleman for expert assistance with liquid chromatography.

## Funding

This work has been financially supported by ReumaNederland (17-1-402 and 08-1-34), the IMI funded project RTCure (777357), ZonMw TOP (91214031) and by Target to B! (LSHM18055-5GF). REMT is a recipient of a European Research Council (ERC) advanced grant (AdG2019-884796). HUS is the recipient of a NWO-ZonMW clinical fellowship (90714509), a NWO-ZonMW VENI grant (91617107), a NWO-ZonMW VIDI grant (09150172010067) and a ZonMW Enabling Technology Hotels grant (435002030) and received support from the Dutch Arthritis Foundation (15-2-402 and 18-1-205). The work has been further funded by the Swedish Research Council (VR Dnr: 2018-02551), the King Gustaf V’s 80-Year Fund, the King Gustaf V’s and Queen Victoria’s Fund, the Swedish Rheumatism Association and the Canadian Institutes of Health Research (CIHR) grant MOP (77700).

## Author contribution

All authors were involved in drafting the article or revising it critically for important intellectual content, and all authors approved the final version to be published.

## Notes

**Conflict of interest** HUS, TWJH and REMT are mentioned inventors on a patent on ACPA-IgG V-domain glycosylation.

### Competing Interest Statement

HUS, TWJH and REMT are mentioned inventors on a patent on ACPA-IgG V-domain glycosylation.

