## Supplementary Material for "ACPA-IgG variable domain glycosylation increases before the onset of rheumatoid arthritis and stabilizes thereafter; a cross-sectional study encompassing over 1500 samples"

Supplementary materials and methods

**Patient and public involvement** – Individuals were involved in this study by donating blood when attending population surveys [1, 2], medical health check-ups [3, 4], were recruited to take part in the arthralgia study in the Amsterdam area of the Netherlands [5] or the treatment strategy study “Improved” in early RA [6].

**Ethical considerations** - All participants have given their written informed consent and the Regional Ethical Review Board Committees approved the studies.

**Laboratory analyses** – ACPA IgG levels were analysed in serum samples using anti-cyclic citrullinated peptide (CCP) enzyme linked immunoassays as previously described [1-3, 5, 7]. ACPA-IgG levels from longitudinal samples of individuals visiting the Early Arthritis Clinic (EAC) were analyzed using an in house anti-CCP2-IgG ELISA.

Supplementary tables

**Table S1. Descriptive cohort data.** Cohorts including samples from individuals in different disease stages from healthy symptom-free to established RA.

| **cohort** | **cohort 1, Japan (Nagasaki)** | **cohort 2, Canada (Manitoba)** | | **cohort 3, Sweden (Umea)** | | **cohort 4, the Netherlands (Amsterdam, Reade)** | **cohort 5, the Netherlands (Leiden, CSA)** | | **cohort 6, the Netherlands (Leiden, Improved)** | | | |
| --- | --- | --- | --- | --- | --- | --- | --- | --- | --- | --- | --- | --- |
| **disease stage** | **healthy, symptom-free** | **healthy** | **RA- onset** | **pre-symptomatic** | **RA- onset** | **arthralgia** | **arthralgia** | **RA- onset** | **RA- onset** | **4 months after RA- onset** | **8 months after RA- onset** | **12 months after RA- onset** |
|  | **N=58** | **N=126** | **N=23** | **N=228** | **N=126** | **N=239** | **N=38** | **N=26** | **N=130** | **N=117** | **N=112** | **N=117** |
| **sex (female), n(%)** | 38 (66) | 90 (72) | 18 (79) | 145 (64) | 78 (61) | 185 (77) | 29 (76.3) | 22 (84.6) | 88 (67.7) | 79 (67.5) | 78 (69.6) | 78 (66.7) |
| **age (years), mean (SD)** | 66.7 (9.9) | 36 (13.4) | 39 (13.5) | 52.2 (9.4) | 59.7 (9.3) | 48.3 (11.6) | 46.5 (12.5) | 47.0 (11.5) | 51.1 (12.5) | 50.6 (12.8) | 50.7 (12.4) | 51.0 (12.4) |
| **arthritis/RA at follow up, n (%)** | 9 (15.5) | 57 (42) | - | 228 (100) | - | 137 (57.3) | 26 (68.4) | - | - | - | - | - |
| **ACPA positive, n (%)** | 58 (100) | 105 (83.3) | 21 (91.3) | 168 (73.7) | 125 (98.4) | 239 (100) | 33 (86.8) | 21 (80.8) | 130 (100) | 117 (100) | 112 (100) | 117 (100) |
| **VDG positive, n (%)** | 48 (83.8) | 42 (33.3) | 19 (82.6) | 105 (46.1) | 116 (92.1) | 211 (87.9) | 27 (71.1) | 18 (69.2) | 117 (90) | 78 (66.7) | 86 (76.8) | 98 (83.8) |
| **VDG%, median (IQR)** | 58.1 (35.6) | 53.1 (68.3) | 109.9 (48.9) | 97.4 (53.5) | 94.2 (50.8) | 75.3 (48.9) | 70.4 (28.8) | 59.1 (49.1) | 96 (48.2) | 95.9 (45.1) | 101.7 (50.3) | 105.2 (48.1) |

**Table S2. Descriptive cohort data.** EAC cohort samples including individuals that achieve drug free remission (DFR), sustained DFR (DFSR) and individuals with late disease flares.

|  | **cohort 7, the Netherlands (Leiden, EAC)** | | | | | |
| --- | --- | --- | --- | --- | --- | --- |
|  | **RA- onset, do not achieve DFR** | **RA- onset, achieve DFR** | **pre-remission** | **DRF** | **DFSR** | **DFR with late flares** |
|  | **N= 59** | **N=36** | **N=52** | **N=41** | **N=35** | **N=11** |
| **sex (female), n(%)** | 42 (71.2) | 36 (65.5) | | | | |
| **age (years) at RA- onset, mean (SD)** | 49.7 (14.5) | 50.9 (13.8) | | | | |
| **ACPA positive, n (%)** | 59 (100) | 32 (89) | 39 (75) | 33 (80) | 31 (89) | 10 (91) |
| **ACPA levels (AU/mL), median (IQR)** | 7340 (5984) | 1816 (7127) | 3583 (5302) | 3010 (8975) | 2725 (7319) | 4259 (10709) |
| **VDG positive, n (%)** | 59 (100) | 19 (95) | 37 (71.2) | 20 (66.7) | 13 (54.2) | 7 (77.8) |
| **VDG%, median (IQR)** | 83.8 (46) | 57.9 (35.8) | 74.05 (30) | 67.7 (41.5) | 80.15 (37.4) | 78.3 (26.9) |

**Table S3. Association between the recognition of multiple citrullinated epitopes (dependent variable) and VDG or ACPA levels (independent variable) in individuals with arthralgia (*Netherlands, Reade*).**

|  | **B (95% CI)**  **univariable (simple)** | **p value** |
| --- | --- | --- |
| **VDG** | 0.03 (0.02 – 0.04) | <0.001 |
| **log ACPA levels** | 0.84 (0.67 – 1.02) | <0.001 |

**Table S4. Association between the recognition of multiple citrullinated epitopes (dependent variable) and VDG or ACPA levels (independent variable) of individuals at RA- onset and in established RA (*Netherlands, Improved*).**

|  | **B (95% CI)**  **univariable (simple)** | **p value** |
| --- | --- | --- |
| **baseline (ordinal regression analysis)** | | |
| **VDG** | 0.142 (0.0045 – 0.0239) | 0.004 |
| **ACPA levels** | 0.0032 (0.0025 – 0.0038) | <0.0001 |
| **over time (GEE)** | | |
| **VDG** | 0.0089 (0.0052 – 0.0125) | <0.0001 |
| **ACPA levels** | 0.0020 (0.0017 – 0.0022) | <0.0001 |

**Table S5. Changes in ACPA-IgG VDG after RA- onset (*Netherlands, Improved*).**

|  | **VDG (%)** | **p value** | **VDG (%), adjusted^1^** | **p value, adjusted** |
| --- | --- | --- | --- | --- |
| **continues** | | | | |
| **per 4 months** | 2.85 (0.51 – 5.19) | 0.017 | 1.29 (-1.30 – 3.89) | 0.32 |
| **per month** | 0.71 (0.13 – 1.30) | 0.017 | 0.32 (-0.33– 0.97) | 0.048 |
| **per visit** | | | | |
| **RA- onset** | 1 (ref) | - | 1 (ref) | - |
| **4 months** | -3.83 (-10.79 – 3.12) | 0.28 | -0.06 (-8.76 – 8.87) | 0.99 |
| **8 months** | 4.15 (-2.86 – 11.16) | 0.25 | 1.51 (-7.50 – 10.51) | 0.74 |
| **12 months** | 7.49 (0.47 – 14.52) | 0.037 | 3.76 (-4.19 – 11.71) | 0.35 |

^1^ covariates: age, gender, CRP and ACPA levels

**Table S6. Generalized estimating equation (GEE) analysis of VDG from individuals sampled 4 vs. 8 months and 8 vs. 12 months after RA- onset (*Netherlands, Improved*).** Association between VDG and treatment after 4 or 8 months respectively. The regression coefficient (B, with 95% CI) indicates changes in VDG between 4 vs. 8 months and 8 vs. 12 months for the different treatment groups.

|  | **B (95% CI)** | **p value^1^**  **univariable (simple)** |
| --- | --- | --- |
| **VDG (%) 4 vs. 8 months** | | |
| **early remission^2^** | 7.52 (-6.79 – 21.83) | - |
| **treatment escalation^3^** | 3.27 (-6.35 – 12.90) | 0.98 |
| **VDG (%) 8 vs. 12 months** | | |
| **early remission, drug-free^4^** | 12.27 (-7.32 – 31.87) | - |
| **early remission no drug-free and treatment escalation^5^** | 6.42 (-0.35 – 13.19) | 0.007 |

^1^ interaction term *treatment decision*time*

^2^ methotrexate (MTX) monotherapy

^3^ MTX, prednisone, hydroxychloroquine, and sulphasalazine combination (arm 1) or MTX and adalimumab combination (arm 2)

^4^ MTX tapering for individuals achieving DFR

^5^ continuous MTX and prednisone or adalimumab treatment

**Table S7. Logistic regression analysis of individuals at RA- onset (*Netherlands, Improved*).** Association between VDG (dependent variable) at RA- onset and treatment response - remission at 4 months and early drug-free remission within 1 year (independent variable).

|  | **OR (95% CI)** | **p value** |
| --- | --- | --- |
| **remission at 4 months** | 1.003 (0.991 – 1.014) | 0.65 |
| **drug free remission in the first year** | 1.00 (0.99 – 1.01) | 0.50 |

Supplementary figure legends

**Figure S1: ACPA-IgG capturing and liquid chromatography VDG analysis.** (a) Schematic illustration of the experimental procedure. ACPA were captured using CCP2- streptavidin beads, followed by IgG capturing using FcXl beads. *N*-linked glycans were released from ACPA-IgG using PNGaseF and 2-AA labeled for liquid chromatography (UHPLC). Glycan peaks including G0F, G1F, G2F, G2FBS1, G2FS2 and G2FBS2 are highlighted. Agalactosylated (G0), monogalactosylated (G1), digalactosylated (G2), fucose attached to the core GlcNAc (F), bisecting GlcNAc (B), monosialylated (S1), disialylated (S2). Blue square: *N-*acetylglucosamine (GlcNAc), green circle: mannose, yellow circle: galactose, red triangle: fucose, pink diamond: *N-*acetylneuraminic acid. (b) Schematic illustration of the formula used to calculate the percentage of ACPA-IgG VDG. (c) UHPLC chromatograms of ACPA-IgG *N*-linked glycan from a healthy individual (Japan), a subject with arthralgia (the Netherlands, Amsterdam, Reade), and an individual at RA- onset and 4, 8 and 12 months after disease development (Improved). (d) Matched paired ACPA-IgG VDG of individuals diagnosed with RA later in life and sampled prior to symptom-onset and at RA- onset. Data are presented as box and whiskers including all data points. ACPA-IgG VDG show slight, non-significant increase when comparing both groups without specifying different time points pre-disease.

**Figure S2: ACPA-IgG VDG percentages correlate with ACPA levels.** Pearson correlation between ACPA-IgG VDG percentages and ACPA levels (aU/ml). A strong correlation was observed in healthy and arthralgia individuals, moderate associations in pre-symptomatic individuals as well as 4/ 8 months after RA- onset and weaker associations at RA- onset and 12 months thereafter. The respective p-values (two-tailed) and correlation coefficients are presented in the figures.

**Figure S3: ACPA-IgG VDG in individuals with arthralgia (*Netherlands, CSA*).** (a) Longitudinal analysis of ACPA-IgG VDG percentages from individuals with clinically suspect arthralgia (CSA), at RA- onset as well as 1 or 2 years after disease development. (b) Time line of VDG percentages up to 36 months after 1^st^ visit. Time point of RA- onset is depicted in orange.

**Figure S4: Longitudinal analysis of ACPA-IgG VDG at RA- onset and in established RA (*Netherlands, Improved*) separated for the different treatment arms.** (a) ACPA-IgG VDG comparison of the four treatment groups: early remission, drug-free; early remission, no drug-free; arm 1 and arm 2. Treatment specifications are illustrated in figure 3c. (b) Longitudinal matched paired samples are shown for treatment arm 1 and 2. ACPA-IgG VDG are reduced in treatment arm 1, 4 months after RA- onset and show an increase after 12 months. No major changes were found for treatment arm 2.

**Figure S5: ACPA-IgG VDG over various time-points and for different ACPA titer.** (a) Time-line of ACPA-IgG VDG after RA- onset and towards drug-free remission (DFR) (*Netherlands, EAC*). ACPA-IgG VDG mean and SEM of the samples presented in figure 4d is depicted. Individuals that flare later in time are shown in red and show slightly higher VDG percentages over time compared to individuals that stay in DFR (turquoise). (b) Stable VDG of ACPA-IgG positive patient serum measured with different titers. ACPA-IgG titers were lowered by mixing the sera with ACPA negative healthy donor sera. Two individual experiments are presented.
