## Supplementary figures and images for "ACPA-IgG variable domain glycosylation increases before the onset of rheumatoid arthritis and stabilizes thereafter; a cross-sectional study encompassing over 1500 samples"

### Figure S1

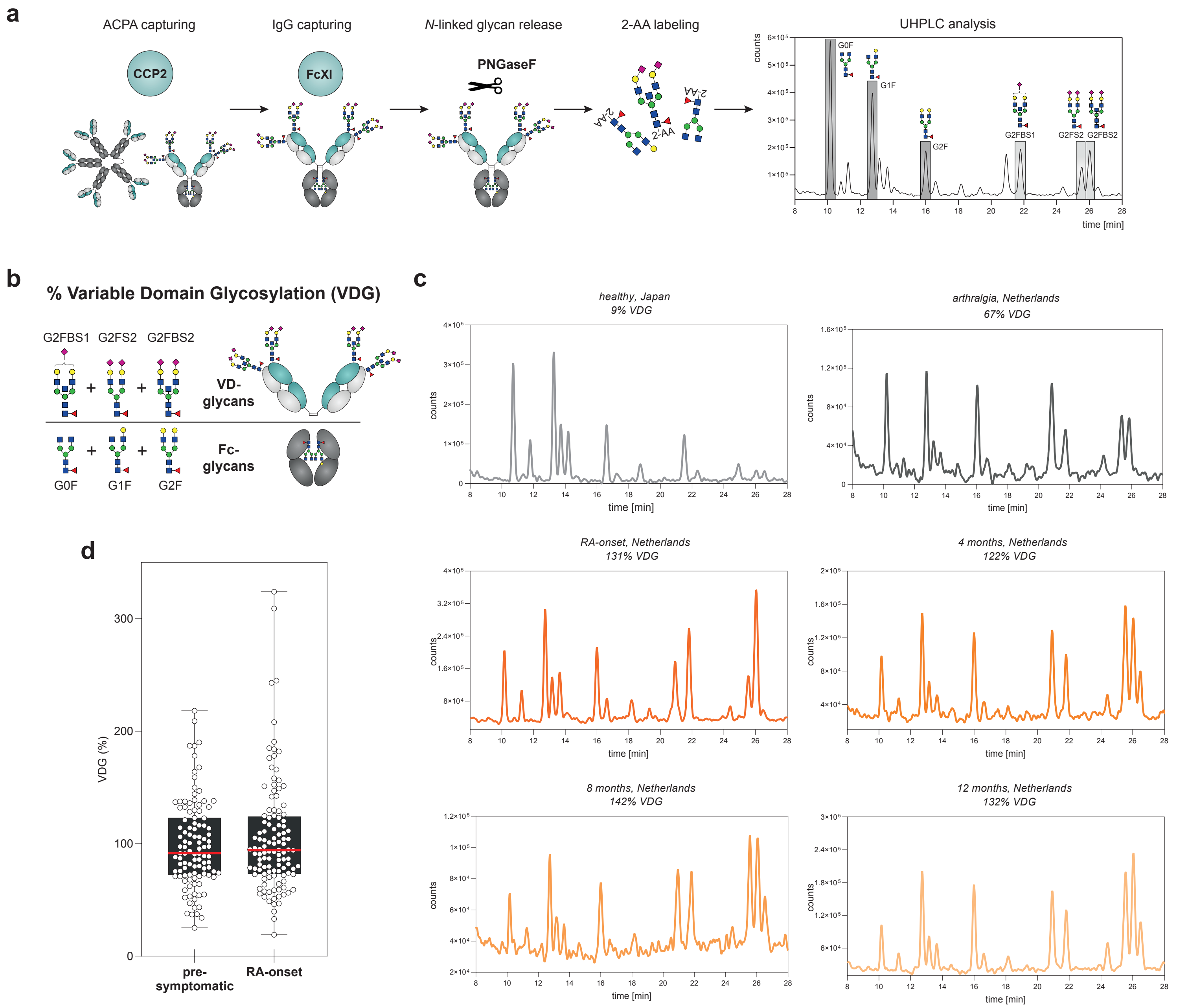

### Figure S2

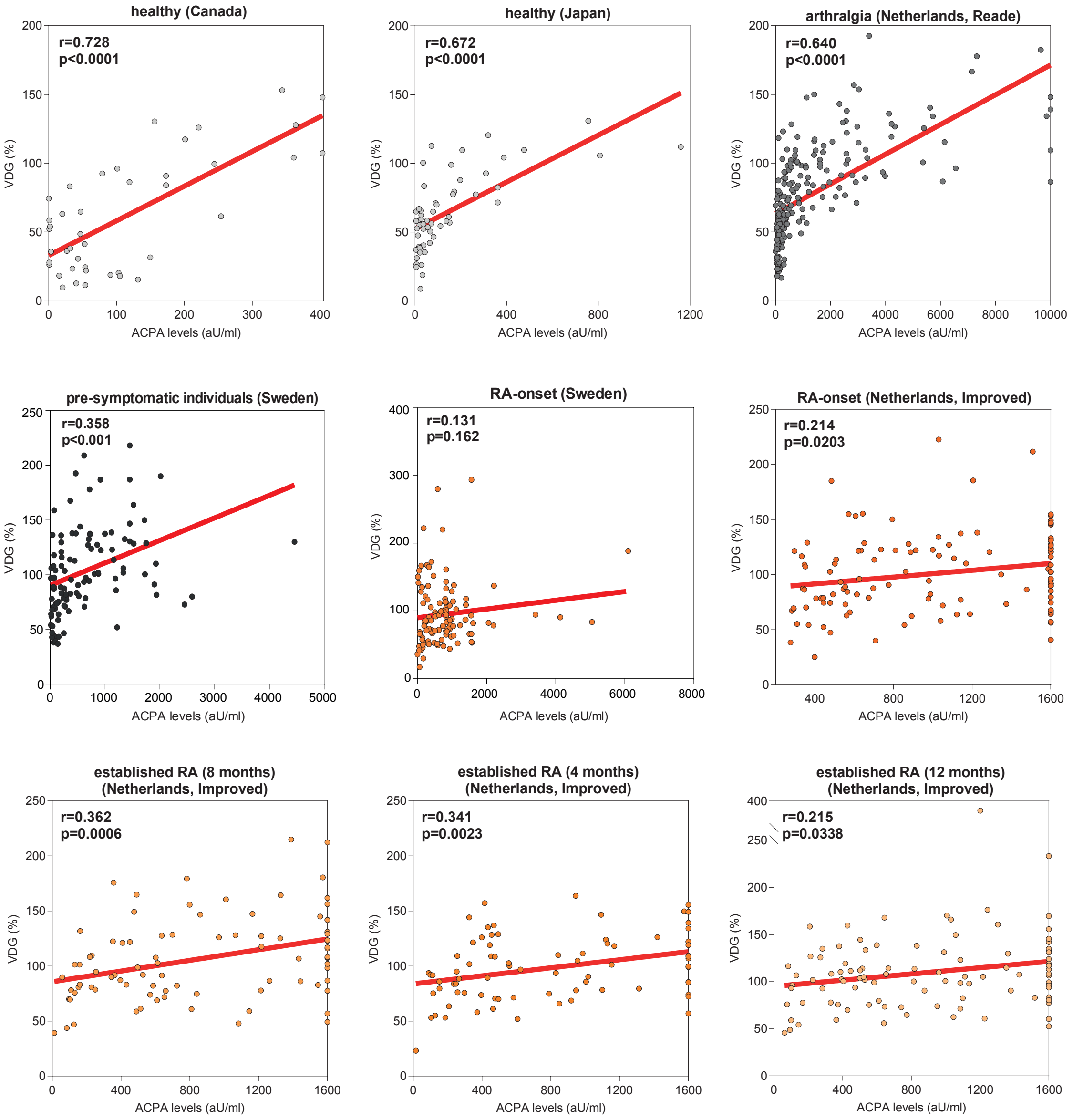

### Figure S3

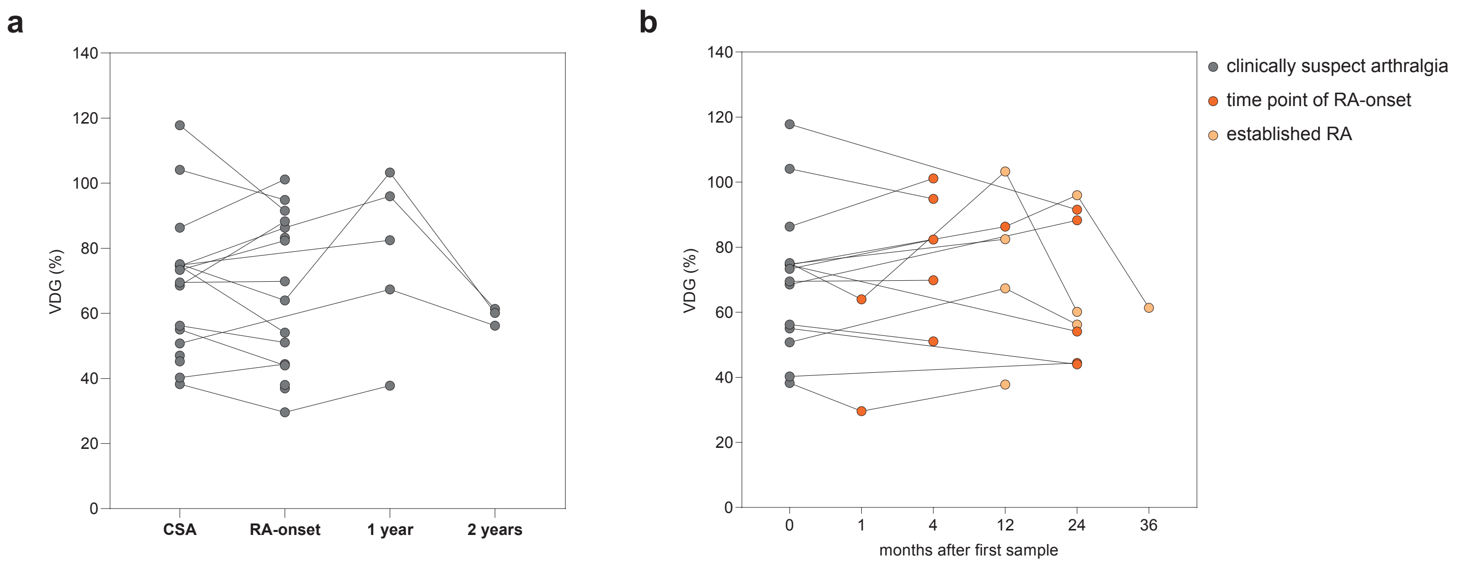

### Figure S4

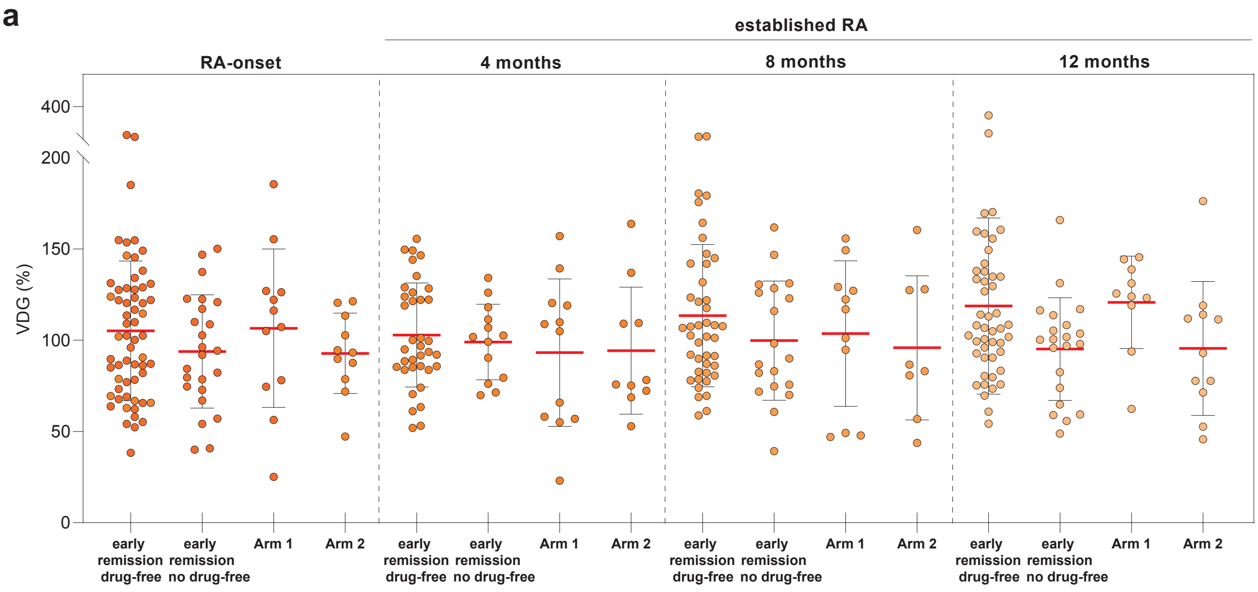

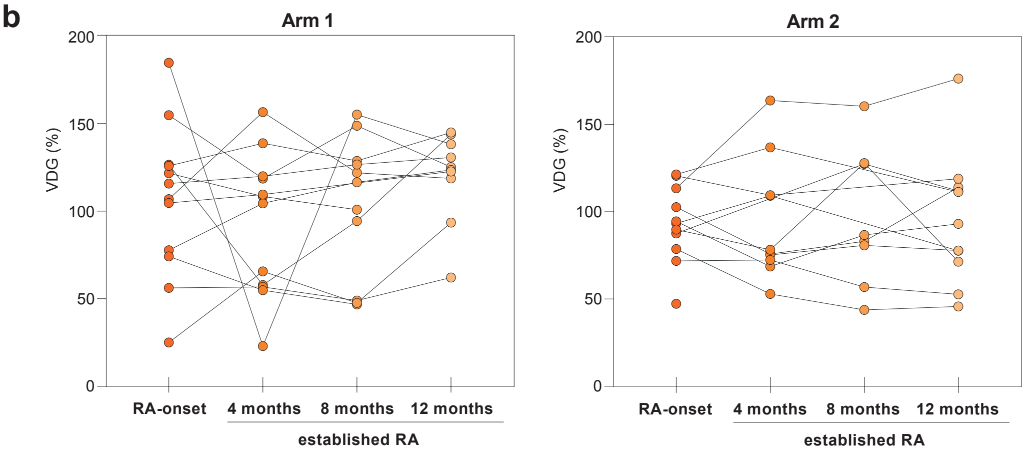

### Figure S5

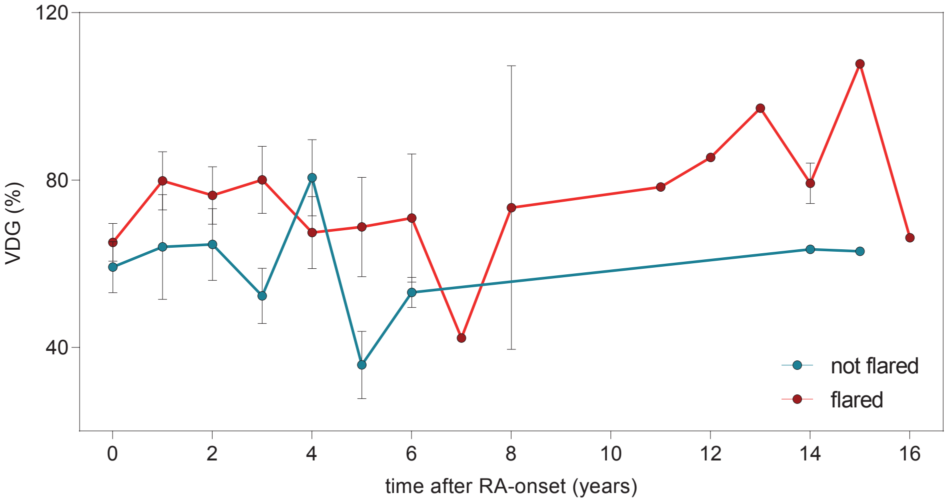

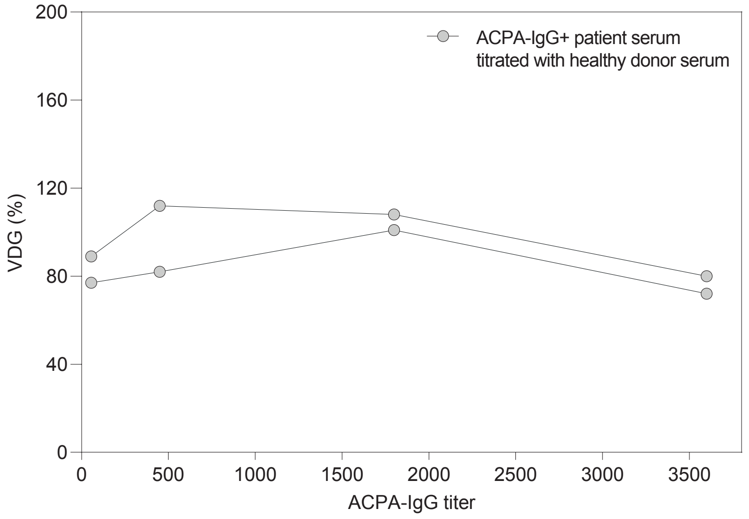
